# A fish finder for birds: route-dependent foraging behavior of the brown booby (*Sula leucogaster*) on a passenger ferry

**DOI:** 10.1101/2025.09.19.676425

**Authors:** Ryota Hayashi

**Affiliations:** Research & Development Center, Nippon Koei Co., Ltd., Tsukuba, Ibaraki, Japan

**Keywords:** Simple observation method, brown booby, Ogasawara Islands, vessel-following behavior, foraging ecology, bottom-up citizen science

## Abstract

This manuscript reports a novel case of route-dependent foraging behavior in the brown booby (*Sula leucogaster*), based on visual observations from a passenger ferry between Chichijima and Hahajima Islands, Ogasawara, Japan. On four dates in 2017, the number of accompanying boobies and their plunge-diving behavior were recorded using only handheld counters. More birds and dives were observed on routes departing from the nesting-rich Hahajima, with peak activity mid-route. These findings suggest that the ferry acts as a “fish finder,” flushing prey such as flying fish for the birds to exploit. The study highlights how low-tech observations from public transport can provide valuable behavioral insights and underscores the potential of citizen science in seabird ecology.

## Introduction

*“Whales are found in the sea, also huge crawfish, enormous shells, and echini, which are called ‘gall of the sea. The ocean here is unusually rich in various products*.*”* (Hawks, 1856)

A journey aboard the local ferry *Hahajima-Maru*, which connects the islands of Chichijima and Hahajima in Japan’s Ogasawara archipelago, known as Bonin Islands, often provides a spectacle of nature as reported by Commodore Perry (Hawks, 1856). As I finished my 24-hour voyage on the *Ogasawara-Maru* and transferred to the *Hahajima-Maru* at Chichijima Island to head to Hahajima Island, I noticed large, conspicuous birds accompanying the ferry throughout the cruise. The birds are the brown boobies, *Sula leucogaster* (Boddaert, 1783), the most conspicuous seabirds in Ogasawara Islands (Ozawa & Saotome, 1968; Saotome, 1968; Yoda et al., 2004). Boobies known for plunge diving (Fig. 1), distribute a widespread tropical area. While many studies have documented seabird associations with commercial fishing vessels that offer scavenging opportunities (e.g. Cherel et al. 1996; Votier et al. 2010; Yorio et al. 2010), interactions with non-fishing vessels—such as passenger ferries—remain poorly studied, as these ferries, unlike fishing vessels, do not provide scavenging opportunities. This study was conducted in the Ogasawara Islands, primarily breeding area for the brown boobies, a subtropical oceanic archipelago located approximately 1,000 km south of mainland Japan. Major colonies are concentrated on uninhabited islets surrounding Hahajima Island (Chiba et al. 2007; Kawaguchi et al. 2014). In contrast, Chichijima Island, the administrative center of the archipelago, hosts very few breeding individuals due to habitat loss and introduced predators such as feral cats (Horikoshi 2007; Kawakami & Masuko, 2008). Despite this asymmetric distribution, brown boobies are frequently observed following the regular inter-island ferry *Hahajima-Maru*, which travels once or twice daily between Chichijima and Hahajima. To document this behavior, visual observations were carried out aboard the *Hahajima-Maru* (Supplementary Movie S1), and the number of accompanying brown boobies and their feeding dives were recorded during four ferry cruises in 2017 (Fig.2). Each one-way trip takes approximately two hours. Brown boobies frequently follow this vessel, which occasionally disturbs flying fish (*Cypselurus* spp.) near the surface during its transit. Visual surveys were conducted during four daytime ferry trips in 2017: April 9, May 12, August 3, and August 10. Two trips were outbound (Chichijima to Hahajima) and two were return trips. Observations were made from both the lower and upper decks of the ferry to maximize the field of view. The number of accompanying brown boobies and the number of plunge dives were recorded once per minute using a stopwatch and field notebook. Each observation session began immediately after departure and continued until the ferry reached its destination. GPS tracks of each cruise were recorded using the Runkeeper mobile app (FitnessKeeper Inc., USA) on an Android smartphone to confirm cruise duration and direction. Weather and visibility conditions were noted to ensure survey consistency; all observations were made under clear or partly cloudy conditions with good visibility.

**Fig. 1.**
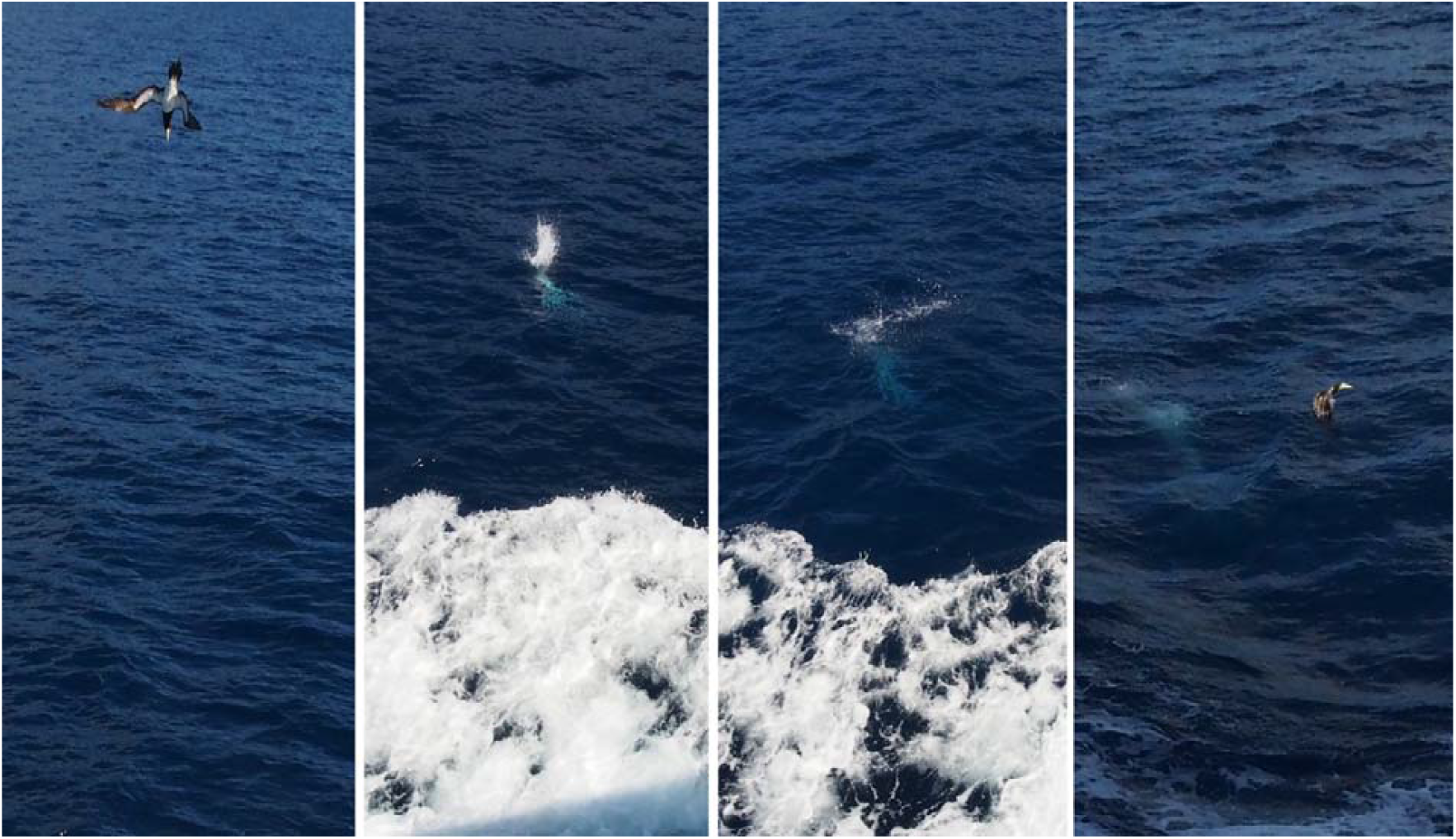
Feeding dive of a brown booby (Sula leucogaster) photographed from the upper deck of the inter-island ferry Hahajima-Maru near Hahajima Island, Ogasawara Islands, Japan. Photographs by Ryota Hayashi.

**Fig. 2.**
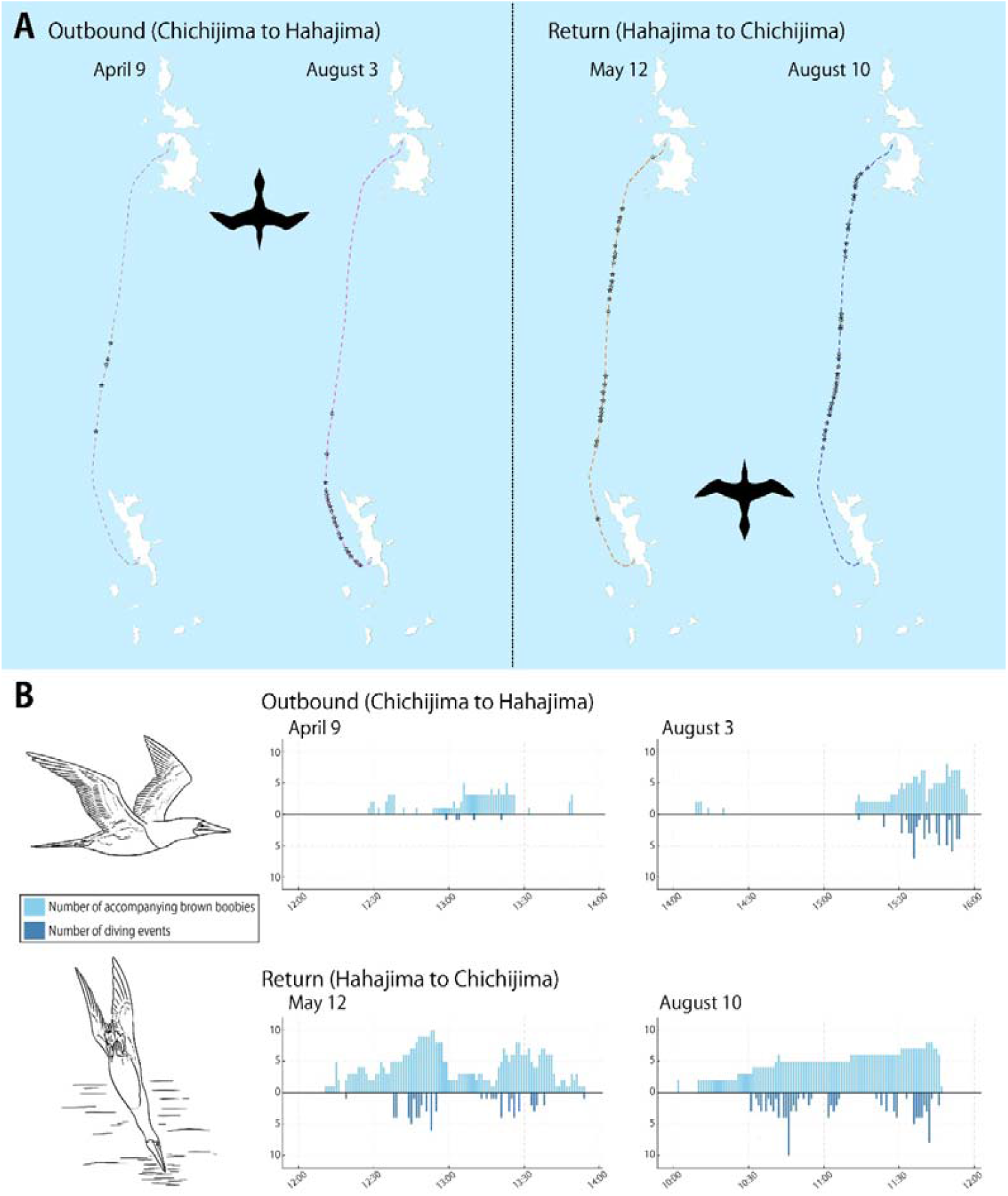
Route-dependent foraging behavior of brown boobies observed from the inter-island ferry Hahajima-Maru. (A) Map showing the ferry routes for the four cruises. Silhouettes indicate the direction of travel for outbound (Chichijima to Hahajima) and return (Hahajima to Chichijima) cruises. Stars mark the locations of feeding dive events. (B) Frequency distribution of the number of accompanying brown boobies (upper panel) and feeding dive events (lower panel) for each cruise, plotted along a timeline from departure to arrival. Map design and silhouettes by Ryota Hayashi; booby flying and diving illustrations by Dr. Chihiro Kinoshita, used with permission.

### Obserbation

Brown boobies were observed accompanying the ferry on all four cruises. However, a consistent directional pattern was found in both the number of individuals and the frequency of feeding dives (Fig. 2). On the two outbound cruises from Chichijima to Hahajima (April 9 and August 3), boobies were rarely present in the first half of the route, boobies began to appear approximately halfway, and increased in number as the vessel approached Hahajima. The closely related species, northern gannet *Morus bassanus* (Linnaeus, 1758), is possible to find a fishing vessel for scavenging feeding about 11 kilometers ahead (Bodey et al. 2014). This suggests that the brown boobies around Hahajima Is. could not find *Hahajima-Maru* before the half way point in the forward cruise. In contrast, on the two return cruises from Hahajima to Chichijima (May 12 and August 10), brown boobies were already present immediately after departure and remained active throughout the route. The total number of accompanying individuals per minute was substantially higher on return trips (431 and 474) compared to outbound trips (108 and 193). Similarly, the number of feeding dives was markedly greater on return trips (71 and 113) than on outbound trips (5 and 59). Notably, brown boobies’ activity appeared higher in August than in April and May; for instance, the return cruise in August recorded 113 dives, compared to 71 in May. The higher activity observed in August coincides with the peak chick-rearing period for brown boobies in the Ogasawara Islands (Momiyama, 1930) and it takes about three months from hatching to leaving the nest (Ospina-Alvarez, 2014). This suggests that parents may rely more heavily on this predictable food source when provisioning hungry offspring. In all cruises, feeding dive frequency increased near the presumed breeding grounds around Hahajima. These findings suggest that presence and foraging activity of brown boobies are spatially structured and may be influenced by colony proximity. While the limited number of cruises (n=4) precludes formal statistical analysis, the consistency of this pattern across multiple dates and directions strongly supports its biological relevance.

## Discussion

This feeding behavior using the regular inter-island ferry is particularly fascinating given the species’ Japanese name, ‘Katsuodori’, which translates to ‘skipjack bird’. This name originates from its traditional association with schools of skipjack tuna (*Katsuwonus* spp.) that drive prey to the surface—a phenomenon historically used by fishermen as a natural fish finder. This report suggests a remarkable role reversal: the Katsuodori now appear to be using the ferry as their own ‘fish finder’ to exploit prey startled by the vessel.

This discovery opens up numerous questions regarding individual variation and social learning within the population. For instance, what type of individuals exploit the ferry? Are they subordinate individuals outcompeted during prime foraging times, or are they innovative individuals that have discovered an efficient strategy? Furthermore, how is this behavior transmitted? Is it a culturally transmitted tradition, perhaps passed from parent to offspring, or does it spread through other social learning mechanisms? These questions lead directly to deeper inquiries about animal cognition. The consistent, spatially structured pattern suggests that brown boobies may possess sophisticated cognitive abilities, such as a cognitive map of the area and associative learning. This interaction also prompts intriguing questions about the boundaries of animal tool use (Seed & Byrne, 2010). Answering these questions requires moving beyond simple observation. Future studies integrating bio-logging technologies (Watanabe & Papastamatiou, 2023; Sato et al. 2025), such as GPS trackers combined with accelerometers and animal-borne video loggers, would be invaluable. Such tools could help identify which individuals use the ferry, quantify their foraging success relative to others, and potentially reveal the cognitive mechanisms and learning processes underlying this remarkable adaptation.

Importantly, this study demonstrates that meaningful behavioral data can be collected using a simple, low-cost method requiring no specialized equipment. In this study, I used only a pen, note, watch, and GPS app installed to Android smartphone, there is no need any specialized devices, however, author could not show here sufficient trials of survey unfortunately. It is because the trip of Ogasawara Islands is limited by the local regular lines, and one-way needs 24 hours to Chichijima and more 2 hours to Hahajima Is. Anyone can survey without needing any special equipment as long as they are aboard the *Hahajima-Maru* and Tourists using *Hahajima-Maru* also conduct this visually survey easily. If the survey results by tourists will be accumulated in future, it will contribute the advances of animal ecology and cognition. Ogasawara Islands are attractive subtropical area, 32577 tourists had come in 2017, however, Ogasawara is extremely far to investigate easily for non-islanders. Recently, the citizen science has been focused in the ecological studies, but many of them were used for biodiversity conservation by monitoring by many eyes. This accessible approach is easily replicable and scalable, offering a powerful tool for ecological monitoring, particularly in remote regions. Its simplicity also makes it ideal for citizen science initiatives. While most citizen science projects follow a top-down, researcher-driven “Contributory model,” the method proposed here enables bottom-up, “Co-created model” research that begins with the curiosity and discoveries of citizens themselves (Haklay 2012; Shirk et al. 2012). This case study suggests that anyone, even non-experts, can become a bearer of scientific discovery with simple tools and a keen eye on their environment.

## ACKNOWKEDGEMENTS

This study was funded by the JSPS KAKENHI Grant Numbers JP16K21005 to RH. The illustrations of brown boobies in Figure 2 were made by Dr. Chihiro Kinoshita.

## CONFLICT OF INTEREST STA TEMENT

The author declares no conflicts of interest.

## Open Research statement

Data is included in our video file.

